# VS-Suite: Automated Consensus Virtual Screening Platform for Improved Screening of Small Molecules

**DOI:** 10.64898/2026.11.17.766942

**Authors:** Ghate Purohith Kumar, N Giri Lakshman, S Devikala, Venkatesan Subramanian

## Abstract

Growing the recent advances in artificial intelligence and machine leaning (AI&ML) approaches accelerated small molecules screening against targeted proteins with limited computational resources and time. Finding accurate binding pose and affinity prediction is challenges task and traditional docking tools and lacks the reproducibility. Development of the Co-folding model is paradigm shift in drug discovery and development, allowing to predict protein ligand complexes without experimental crystal structures. Notably Boltz-2 predicts the complex structure along with binding affinity directly using protein FASTA sequence and ligand SMILES. Our previous studies confirmed that Boltz-2 is classify binder non-binders effectively. Autodock Vina and GNINA are widely used docking with knowledge and CNN based scoring functions allows to screen potential small molecules. We developed in-house VS-suite (Virtual Screening Suite) multi-environment conda orchestration designed to automate the docking workflow only using FASTA and SMILES string as inputs via streamlit-based browser GUI which integrates the boltz-2 for the complex structure prediction followed docking using AutoDock Vina for Gnina. Validation on 58 antiviral nucleoside analogues screened against SARS-CoV-2 main protease (Mpro). VS-Suite is available at https://github.com/Vsubuchem-lab/Virtual-screening-suite-VS-Suite. This consensus approaches helpful in rapid screening of small molecules against selected targets. The findings and VS-suite support its utility as a rapid and reliable tool for hit identification and rational drug development.

## Introduction

Many drugs used in the treatment of various diseases can be traced back to natural compounds derived from medicinal herbs and traditional medicinal plants. Throughout human history, medicinal plants have played a significant role as valuable sources of therapeutic agents. Various plant parts, including leaves, roots, bark, fruits, and seeds, have been traditionally utilized for their diverse medicinal properties and therapeutic benefits. Cragg et al have extensively documented the significance of natural products in modern drug discovery. They have emphasized the importance of medicinal plants and other natural sources as rich reservoirs of bioactive compounds and potential lead molecules for pharmaceutical research and drug development. In parallel, advances in computational infrastructure, particularly the increasing processing capabilities of modern hardware and the development of sophisticated algorithms and molecular modelling software packages have significantly revolutionized virtual screening approaches, enabling more efficient and effective identification of potential drug candidates.

Early molecular docking algorithms primarily focused on rigid-body geometric fitting and shape complementarity. Subsequent advancements introduced ligand flexibility, followed by the incorporation of protein flexibility through flexible docking and induced-fit docking approaches. Improvements in conformational sampling techniques have further enhanced the ability to predict protein–ligand interactions and estimate binding affinities with greater accuracy. More recently, the integration of GPU-accelerated computing, large-scale chemical databases, machine learning, and deep learning approaches has significantly expanded the computational efficiency, scalability, and predictive capabilities of molecular docking, enabling high-throughput screening of increasingly large chemical libraries for potential lead compounds.

Conventionally, molecular docking is widely employed in computational drug discovery to screen potential drug candidates against specific protein targets.□□□□□ However, more rigorous estimation of binding free energies using molecular dynamics (MD) simulation trajectories, including endpoint free energy methods such as molecular mechanics Poisson–Boltzmann surface area (MM-PBSA) and molecular mechanics generalized Born surface area (MM-GBSA), enhanced sampling techniques such as umbrella sampling and metadynamics, and free energy perturbation (FEP), is computationally demanding and time-consuming. Although these methods provide more reliable estimates of protein–ligand binding free energies, their high computational cost generally limits their application to a relatively small set of selected molecules.

### Emerging of computational approaches for drug discovery

The theoretical foundation of molecular recognition can be traced back to the late nineteenth century, when Emil Fischer proposed the lock-and-key model in the 1890s. This model conceptualized the receptor as a rigid structure with a specific binding-site geometry, into which a ligand with complementary shape and chemical properties could fit. Both the receptor and ligand were assumed to be rigid, with no consideration of conformational flexibility. Since then, molecular docking has undergone substantial evolution, progressing from rigid-body geometric matching to flexible, physics-based simulations and, more recently, data-driven approaches based on machine learning and artificial intelligence. Broadly, the evolution of molecular docking can be classified into three generations. First-generation methods were primarily based on the lock- and-key concept, relying on rigid-body docking and geometric shape complementarity. Second-generation methods incorporated ligand conformational flexibility through systematic or stochastic sampling techniques, including genetic algorithms, coupled with empirical and force-field-based scoring functions. Modern, third-generation approaches increasingly leverage machine learning, deep learning, and generative artificial intelligence to improve conformational sampling, binding-pose prediction, and the identification of promising drug candidates

Biological macromolecules can undergo conformational changes to accommodate the structural and chemical characteristics of their binding partners. In 1958, Koshland introduced the induced-fit theory, which proposed that both the ligand and receptor can undergo conformational adjustments upon binding to achieve optimal molecular complementarity. The development of computational approaches for predicting receptor–ligand interactions began in the early 1980s. In 1982, Kuntz and colleagues developed UCSF DOCK, one of the earliest computational molecular docking programs, which was primarily based on the lock-and-key concept and relied on rigid-body geometric matching between the receptor and ligand.

Second-generation molecular docking programs incorporated greater conformational flexibility and employed empirical, force-field-based, and knowledge-based scoring functions to improve binding-pose prediction and affinity estimation. Several widely used docking programs emerged during this period, each implementing distinct conformational sampling and optimization strategies. These include AutoDock, which employs stochastic search algorithms; GOLD, which utilizes genetic algorithms; FlexX, which adopts a fragment-based incremental construction approach; Glide, which employs hierarchical filtering and systematic conformational searching; Surflex, which uses surface-based molecular similarity; rDock, which combines stochastic and deterministic search strategies, including genetic algorithms and Monte Carlo methods; AutoDock Vina, which utilizes gradient-based optimization; and GNINA, which integrates gradient-based pose optimization with convolutional neural network (CNN)-based rescoring. The key characteristics and computational strategies of these docking programs are summarized in **Table 1**.

**Table 1:** Historical development of molecular docking tools and their computational approaches.

| SNO | Docking method/tool | Search algorithm | Scoring function | Year of Release |
|---|---|---|---|---|
| 1 | UCSF DOCK | Geometric matching; shape complementarity | Geometric matching; shape complementarity | 1982 |
| 2 | AutoDock 1 | Metropolis Monte Carlo; simulated annealing | Metropolis Monte Carlo; simulated annealing | 1990 |
| 3 | FlexX | Incremental construction; fragment-based placement | Incremental construction; fragment-based placement | 1996 |
| 4 | GOLD | Genetic algorithm | Gold-Score (force-field-based); Chem-Score (empirical) | 1997 |
| 5 | ICM | Biased-probability Monte Carlo; local optimization | Biased-probability Monte Carlo; local optimization | 1994 |
| 6 | FRED | Exhaustive rigid-body search; conformer-based docking | Exhaustive rigid-body search; conformer-based docking | 2003 |
| 7 | Surflex | Molecular similarity-based search; fragment placement and optimization | Molecular similarity-based search; fragment placement and optimization | 2003 |
| 8 | Glide | Systematic conformational, positional and orientational search; hierarchical filtering | Systematic conformational, positional and orientational search; hierarchical filtering | 2004 |
| 9 | HADDOCK | Information-driven docking using rigid-body energy minimization, semi-flexible simulated annealing and flexible refinement | HADDOCK score (weighted combination of intermolecular energy, van der Waals, electrostatic, desolvation and restraint terms) | 2005 |
| 10 | PLANTS | Ant Colony Optimization (ACO) | Ant Colony Optimization (ACO) | 2009 |
| 11 | AutoDock 4 | Lamarckian Genetic Algorithm (LGA); Monte Carlo-based local search | Lamarckian Genetic Algorithm (LGA); Monte Carlo-based local search | 2009 |
| 12 | AutoDock Vina | Iterated Local Search (ILS); BFGS local optimization | Iterated Local Search (ILS); BFGS local optimization | 2010 |
| 13 | smina | Vina-derived stochastic search and local optimization | Vina-derived stochastic search and local optimization | 2013 |
| 14 | DOCK 6 | Anchor-and-grow; incremental construction | Anchor-and-grow; incremental construction | 2015 |
| 15 | Prodigy-LIG* | Not applicable (binding-affinity prediction rather than docking) | Not applicable (binding-affinity prediction rather than docking) | 2018 |
| 16 | GNINA | Vina-based Monte Carlo sampling and local optimization | Vina-based Monte Carlo sampling and local optimization | 2021 |

\*Affinity predictor

**Table 1:** Representative VS-Suite results for 58 antiviral compounds screened against SARS-CoV-2 Mpro (in apathetical order).

| **Compound** | **Boltz IC□□ (µM)** | **Boltz P(binder)** | **Boltz ipTM** | **Vina ΔG (kcal/mol)** | **GNINA ΔG (kcal/mol)** | **GNINA CNN pose** | **GNINA CNN pK** |
|---|---|---|---|---|---|---|---|
| **Abacavir** | 34.14 | 0.17 | 0.67 | -8.00 | -7.90 | 0.87 | 5.92 |
| Acyclovir | 144.70 | 0.20 | 0.97 | -5.30 | -4.30 | 0.78 | 4.39 |
| Adafosbuvir | 66.00 | 0.18 | 0.93 | -8.10 | -7.70 | 0.94 | 6.69 |
| Adefovir | 19.09 | 0.26 | 0.83 | -6.30 | -5.95 | 0.53 | 4.50 |
| Alamifovir | 30.24 | 0.12 | 0.77 | -7.40 | -6.80 | 0.88 | 6.65 |
| Alovudine | 3.49 | 0.10 | 0.70 | -6.30 | -5.50 | 0.74 | 4.26 |
| Amantadine | 944.20 | 0.14 | 0.95 | -4.50 | -2.50 | 0.67 | 3.22 |
| Amdoxovir | 4.91 | 0.27 | 0.87 | -7.20 | -6.70 | 0.67 | 4.42 |

**Table 2:** AI based co-folding models for bio-molecular structure predictions.

| **Co-fold models** | **Year of release** |
|---|---|
| AlphaFold Multimer | 2021 |
| U-Mol | 2023 |
| Boltz-1 | 2024 |
| Chai-1 | 2024 |
| AlphaFold2 | 2024 |
| AlphaFold3 | 2024 |
| Rosetta-Fold All-atom | 2024 |
| NeuralPLexer3 | 2024 |
| Boltz-2 | 2025 |
| Chai-2 | 2025 |
| Protenix | 2025 |

These classical scoring functions are widely used in the early stages of hit identification and virtual screening. However, they have several inherent limitations. Their performance relies heavily on experimentally determined ligand-bound crystal structures (holo-structures) and often fails to adequately account for receptor flexibility. Furthermore, their predicted binding affinities frequently exhibit poor correlation with more rigorous physics-based free-energy methods, such as MM/PBSA, MM/GBSA, and free-energy perturbation (FEP), as well as with experimentally determined binding affinities.

### Development of Co-folding models

Over the past decade, biomolecular structure prediction has undergone a significant transformation, shifting from a predominantly experiment-driven field towards AI-based computational approaches. Initially, these methods and tools focused primarily on predicting individual protein structures. However, recent advances have expanded their capabilities to predict and model a wide range of biomolecular complexes, including protein–ligand, protein–protein, protein–antibody, and protein–nucleic acid interactions.

The development of AlphaFold, driven by advances in artificial intelligence (AI), has revolutionized three-dimensional (3D) protein structure prediction, addressing what was long regarded as the “holy grail” of structural biology. In recent years, co-folding approaches have attracted considerable attention due to their ability to predict 3D protein-ligand complex structures directly from primary amino acid sequences and the Simplified Molecular-Input Line-Entry System (SMILES) representations of small molecules. Building on these advances, several AI-based frameworks, including Chai-1, Boltz-1, and Protenix, have adopted co-folding strategies for biomolecular complex structure prediction, leveraging advanced diffusion architectures and training on accessible structural datasets while maintaining high prediction accuracy. Similarly, RoseTTAFold All-Atom (RF-AA) integrates residue-level representations of proteins and nucleic acids with atom-level representations of small molecules to facilitate all-atom biomolecular complex modelling. UMol and NeuralPLexer have further advanced sequence-to-complex prediction through flexible, all-atom generative frameworks. Particularly, Boltz-2 has emerged as a promising approach by extending 3D protein-ligand complex prediction to include the estimation of binding affinities. Several studies have reported its promising performance in discriminating active from inactive compounds, highlighting its potential utility in large-scale virtual screening.

Collectively, these models offer promising approaches for the structural prediction of biomolecular complexes, virtual screening, and molecular design. However, their predictions require independent structural, biophysical, or experimental validation before they can be interpreted as evidence of molecular binding or biological activity.

In this study, we introduce a VS-suite-based consensus docking approach integrated with Boltz-2 for biomolecular complex structure prediction, followed by molecular docking using Auto Dock Vina and GNINA. This approach builds upon our previous studies evaluating the performance of multiple consensus scoring functions. The VS-suite provides automated docking protocols, thereby minimizing user-induced errors, reducing procedural variability, and improving the reproducibility and consistency of the docking workflow.

## Methods

### Virtual Screening Suite (VS-Suite) Architecture

The Virtual Screening Suite (VS-Suite) is a multi-environment Conda orchestration framework developed in Python 3.10. The framework was designed to overcome package and dependency conflicts among Boltz-2, Open Babel, and RDKit, which require different and potentially incompatible library versions when installed within a single environment. To ensure compatibility and maintain software modularity, VS-Suite is organized into three independent Conda environments: (1) **autodock-main**, which contains RDKit, Bio-python, Streamlit, and other supporting utilities; (2) **autodock-prep**, which includes Open Babel 3.1, PDB2PQR 3.6, PROPKA 3.5, fpocket, and Meeko 0.7 for molecular preprocessing and input preparation; and (3) **boltz2**, a dedicated PyTorch environment for Boltz-2 inference. AutoDock Vina (v1.1.2) and GNINA (v1.1) are integrated as precompiled binaries accessible through the system PATH. Inter-environment communication is managed through a subprocess wrapper using conda run, enabling the main environment (autodock-main) to invoke and coordinate operations in the preparation (autodock-prep) and Boltz-2 (boltz2) environments without direct dependency interference. Thus, autodock-main serves as the central orchestration environment, coordinating all computational tools through subprocess calls.

The Streamlit-based browser interface (VS_GUI.py) provides a user-friendly, three-page graphical interface (Figure 1): (i) **New Run**, which allows users to upload input files and configure docking parameters; (ii) **Results Browser**, which facilitates result inspection, three-dimensional visualization of protein–ligand complex poses, and pose downloading; and (iii) **VS-Suite Workflow and Documentation**, which provides an overview of the computational workflow and accompanying documentation.

**Figure 1.**
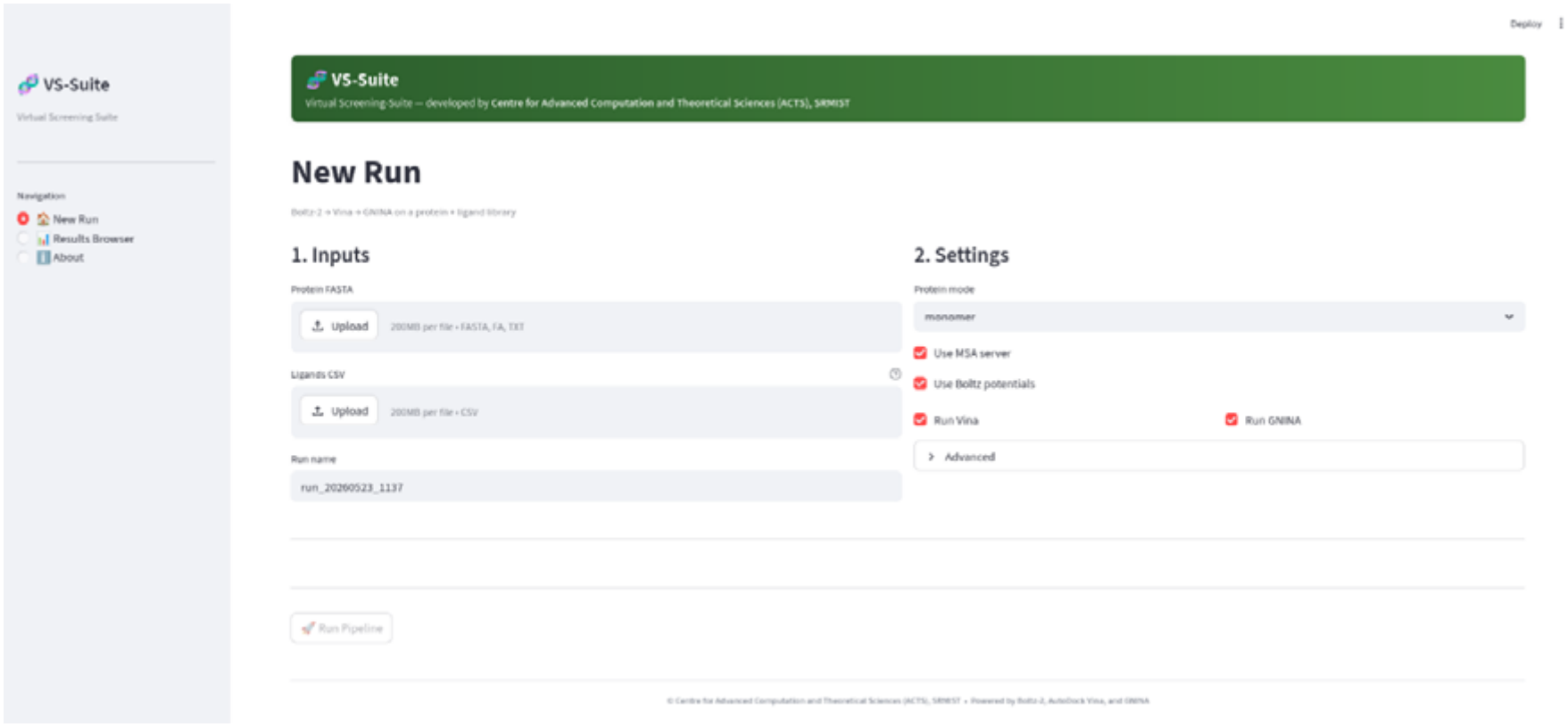
Graphical User Interface (GUI) for the Virtual screening Suite (VS-suite)

**Figure 2.**
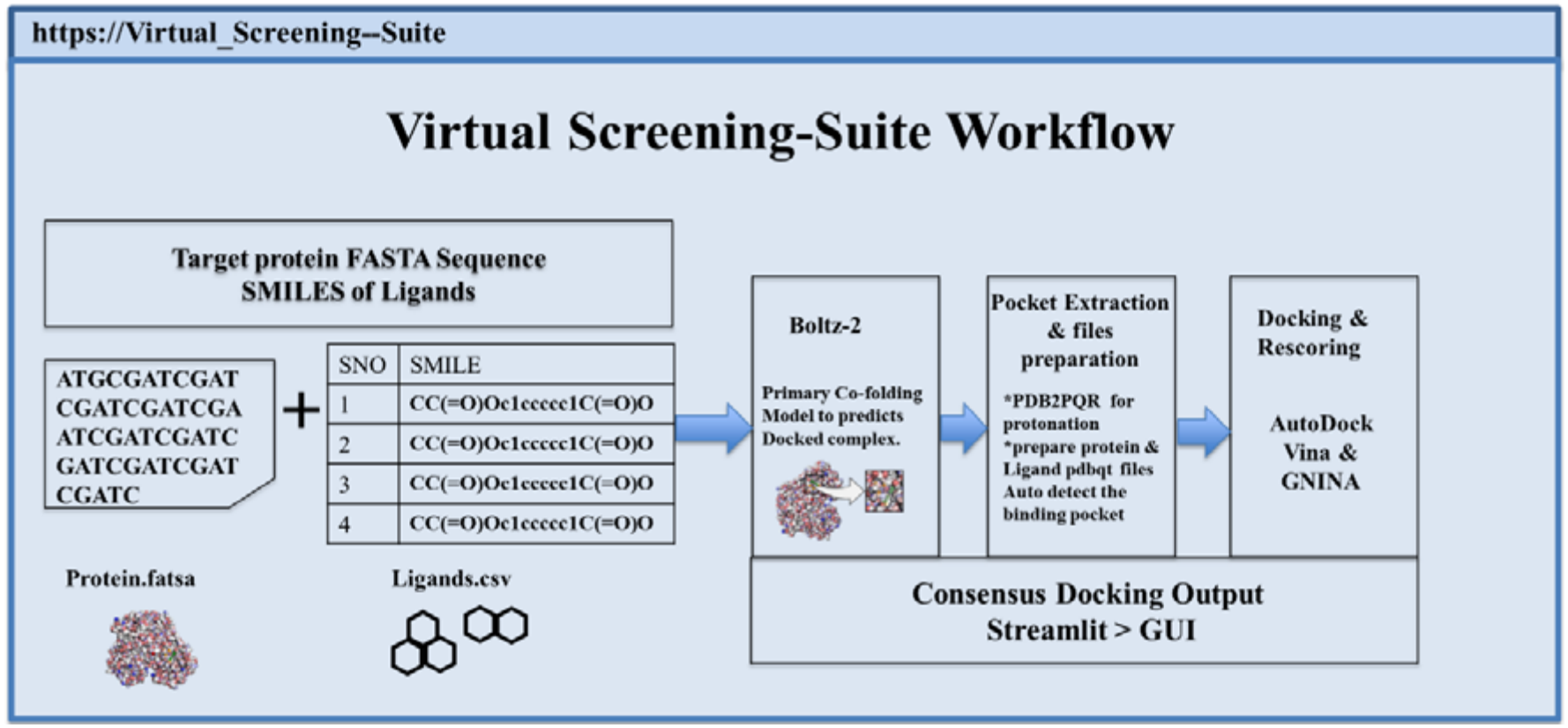
Schematic representation Virtual Screening-Suite (VS-Suite), depicted the steps involved automate the consensus docking.

### Input Preparation for VS-Suite

The targeted protein’s FASTA sequence is encoded into a YAML file to serve as the input for Boltz-2. For subsequent docking, the protein structure is extracted from the Boltz-2 complex, protonated pH 7.4 using PDB2PQR v3.6 with PROPKA pKa estimates, and converted to PDBQT format with Gasteiger partial charges via OpenBabel. The canonical SMILES string serves as ligand inputs provided in CSV file with ligand name and SMILE string. VS-suite supports CSV column headers as ‘name’ or ‘compound’ for ligand name and ‘smiles’ or ‘smi’ for SMILES string. The case insensitive are verified by RDKit, during the YAML generation phase, invalid entries are identified and skipped with warning. Similarly, for Boltz-2 predicted ligand poses were converted to PDBQT using Meeko v0.7, with automatic fallbacks to OpenBabel alternates if Meeko encounter error.

### VS-Suite Workflow

The VS-Suite workflow is executed in four consecutive steps performed for each ligand in the compound library shown in **Figure2**. Step 1, the orchestrator defines the FASTA and CSV inputs for the protein and ligands respectively. For each ligand one Boltz-2 YAML file will be generated. In step 2, Boltz-2 environment invokes and predicts the protein ligand complex as well as affinity. In step 3, binding site is defined from the Boltz-2 output complex structure. The receptor and ligand files are then prepared as mentioned in input preparation, then docking runs using AutoDock Vina first. In the final step, GNINA is executed on the same prepared inputs. The results for each ligand updated in master JSON file.

VS-Suite addresses the practical challenges that limit the adoption the of multiple docking tools. The key aspects of the suite are the systematic integration by orchestrating three complementary scoring functions within single automated workflow. VS-Suite allows cross tool validation for screening of potential compounds targeting protein of interest.

### VS-Suite Performance and Robustness

To quantify the practical performance of VS-Suite, per-stage wall-clock runtimes and completion statuses were recorded during a benchmark run involving 58 natural-product ligands targeting SARS-CoV-2 Mpro on a single NVIDIA RTX A2000 GPU node (12 GB VRAM). All seven pipeline stages, encompassing Boltz-2 structure and binding-affinity prediction, output parsing, receptor and ligand protonation and PDBQT preparation, Auto Dock Vina docking, and GNINA rescoring, were completed successfully for all 58 ligands, yielding a 100% completion rate without requiring any manual intervention. These results demonstrate the robustness of the automated orchestration framework in processing chemically diverse natural-product ligands without requiring ligand-specific parameter tuning or manual adjustments.

## Results and Discussion

VS-Suite addresses a practical barrier that has long hindered the adoption of state-of-the-art docking tools in wet laboratories: the lack of a unified, reproducible, and GUI-accessible pipeline integrating AI-based affinity prediction with conventional molecular docking. The primary contribution of VS-Suite lies not in introducing novel algorithms, but in its architectural integration of complementary methods within a single, automated workflow. This integrated framework enables simultaneous cross-validation of predictions while eliminating the need for multiple independent software installations and bespoke scripting pipelines, thereby making advanced virtual screening approaches more accessible, reproducible, and practical for experimental research laboratories.

All 58 compounds were successfully processed through the complete VS-Suite pipeline without manual intervention, demonstrating the robustness and reliability of the automated preparation workflow. Representative results are presented in Table 1. The complete run required 6.9 h of wall-clock time (24,825 s), corresponding to an average of 421 s (7.0 min) per ligand for complex structure prediction and binding affinity estimation. Boltz-2 inference was the dominant contributor to the overall computational cost, accounting for 66.8% of the total runtime (280.9 ± 15.9 s per ligand) including protein-ligand complex and binding energy estimation, followed by Auto Dock Vina (17.5%; 73.6 ± 85.6 s) and GNINA (13.8%; 57.9 ± 65.2 s). The remaining preparation steps, including protonation, PDBQT generation, and output parsing, collectively accounted for less than 2% of the total runtime.

The runtimes of all three computationally intensive stages exhibited strong positive correlations with ligand heavy-atom count (Spearman’s ρ = 0.82 for Boltz-2, 0.96 for Vina, and 0.97 for GNINA; all p < 0.0001). These correlations provide a quantitative basis for estimating the computational time required to process compound libraries of varying sizes and molecular complexity using heavy-atom count as a readily accessible predictor. On the hardware used in this benchmark, VS-Suite achieved a throughput of approximately 8-9 ligands per hour for compounds of typical natural-product size (11-65 heavy atoms). Notably, GPU-bound Boltz-2 inference, rather than the conventional docking stages, emerged as the primary computational bottleneck, highlighting its role as the principal constraint on scalability in larger virtual-screening campaigns.

The consensus-ranking strategy implemented in VS-Suite represents a deliberate methodological simplification. Assigning equal weights to tool-specific rankings provides a transparent and readily interpretable aggregation approach while avoiding the need for calibration datasets or parameter optimization required by more complex weighted-combination schemes. However, the current implementation is limited to the binding-affinity assessment tier of the pipeline. Integration with downstream free-energy perturbation (FEP) and MM-PB/GBSA calculations is planned for VS-Suite v2.0 to enable more comprehensive evaluation of predicted binding affinities.

Furthermore, the present validation was conducted using a relatively homogeneous chemical class (nucleoside analogues) against a single biological target (Mpro). Broader validation across diverse target classes and chemical spaces is therefore necessary to establish the generalizability and reliability of the VS-Suite consensus-ranking strategy. Ideally, such evaluations should incorporate established benchmarking datasets with experimentally determined binding data, such as DUD-E and LIT-PCBA. Accordingly, the present study should be viewed as a proof-of-concept demonstrating the technical robustness of the automated pipeline and the consistency of predictions across complementary computational tools, rather than as a definitive validation of its performance as a general-purpose virtual-screening filter. Systematic benchmarking across broader chemical and biological spaces is ongoing.

## Conclusions

We present VS-Suite, an open-source virtual screening pipeline that integrates Boltz-2-based AI co-folding, Auto Dock Vina docking, and GNINA-based CNN rescoring into a unified, automated workflow accessible through a browser-based graphical user interface (GUI). When applied to a panel of 58 antiviral nucleoside analogues targeting SARS-CoV-2 main protease (Mpro), VS-Suite successfully processed all compounds through the complete workflow without manual intervention. The results demonstrated strong correlations among the different scoring approaches, supporting the utility of the consensus-ranking strategy underlying the platform’s hit-prioritization framework. By integrating structure prediction molecular docking, and complementary scoring functions, VS-Suite offers a streamlined and reproducible framework for virtual screening. VS-Suite is freely available https://github.com/Vsubuchem-lab/Virtual-screening-suite-VS-Suite and it is designed for straightforward deployment on Linux workstations equipped with NVIDIA GPUs.

## Corresponding Author

**Venkatesan Subramanian**

*Department of Chemistry, SRM Institute of Science and Technology, Kattankulathur, 603 203, Tamil Nadu, India, Center for Advanced Computational and Theoretical Sciences, SRM Institute of Technology, Kattankulathur*, 603 203, *Tamil Nadu, India. E-mail*

## Authors

**Ghate Purohith Kumar**

*Department of Chemistry, SRM Institute of Science and Technology, Kattankulathur, 603 203, Tamil Nadu, India*

**N Giri Lakshman**

*Department of Chemistry, SRM Institute of Science and Technology, Kattankulathur, 603 203, Tamil Nadu, India*

**Devikala S**

*Department of Chemistry, SRM Institute of Science and Technology, Kattankulathur, 603 203, Tamil Nadu, India*

